# Dynamic threat processing

**DOI:** 10.1101/183798

**Authors:** Christian Meyer, Srikanth Padmala, Luiz Pessoa

## Abstract

During real-life situations, multiple factors interact dynamically to determine threat level. In the current functional MRI study involving healthy adult human volunteers, we investigated interactions between proximity, direction (approach vs. retreat), and speed during a dynamic threat-of-shock paradigm. As a measure of threat-evoked physiological arousal, skin conductance responses were recorded during fMRI scanning. Whereas some brain regions tracked individual threat-related factors, others were also sensitive to combinations of these variables. In particular, signals in the anterior insula tracked the interaction between proximity and direction where approach vs. retreat responses were stronger when threat was closer compared to farther. A parallel proximity-by-direction interaction was also observed in physiological skin conductance responses. In the right amygdala, we observed a proximity by direction interaction, but intriguingly in the opposite direction as the anterior insula; retreat vs. approach responses were stronger when threat was closer compared to farther. In the right bed nucleus of the stria terminalis, we observed an effect of threat proximity, whereas in the right periaqueductal gray/midbrain we observed an effect of threat direction and a proximity by direction by speed interaction (the latter was detected in exploratory analyses but not in a voxelwise fashion). Together, our study refines our understanding of the brain mechanisms involved during aversive anticipation in the human brain. Importantly, it emphasizes that threat processing should be understood in a manner that is both context sensitive and dynamic.

## Introduction

Anticipation of aversive events leads to a repertoire of changes in behavioral, physiological, and brain responses that contribute to the handling of the negative consequences of such events. At the same time, abnormalities in aversive anticipatory processing are thought to underlie many mental disorders, such as anxiety and depression (Grupe and Nitschke, 2013; Dillon et al., 2014). Hence, understanding the brain mechanisms of aversive anticipation is important from both basic and clinical standpoints.

In humans, aversive anticipation has been investigated with paradigms in which punctate cues signal an upcoming negative event (Bocker et al., 2001; Jensen et al., 2003; Nitschke et al., 2006; Simmons et al., 2006; Brown et al., 2008), or by blocked manipulations with constant threat level (McMenamin et al., 2014; Vytal et al., 2014). However, during most real-world situations, aversive anticipation changes dynamically over time. An important factor in determining threat level is proximity, as when a prey reacts differently to the presence of a predator when the latter is proximal compared to distant (Figure 1A) (Blanchard and Blanchard, 1990; Blanchard et al., 2011). Other factors involve direction, namely whether threat is approaching vs. retreating (Figure 1B) and speed, reflecting how fast or slow the threat is moving (Fanselow and Lester, 1988). Some studies have taken initial strides at investigating how some of these factors influence brain responses during aversive anticipation. For instance, the contrast of proximal vs. distal threats revealed functional Magnetic Resonance Imaging (fMRI) responses in a host of brain regions, including the anterior insula, midbrain periaqueductal gray (PAG), and bed nucleus of the stria terminalis (BST) (Mobbs et al., 2010; Somerville et al., 2010); evidence for amygdala involvement linked to threat proximity is mixed (Mobbs et al., 2010; Somerville et al., 2010). Similarly, comparison of approaching vs. retreating threats has revealed responses in the anterior insula, BST, and amygdala (Mobbs et al., 2010).

**Figure 1.**
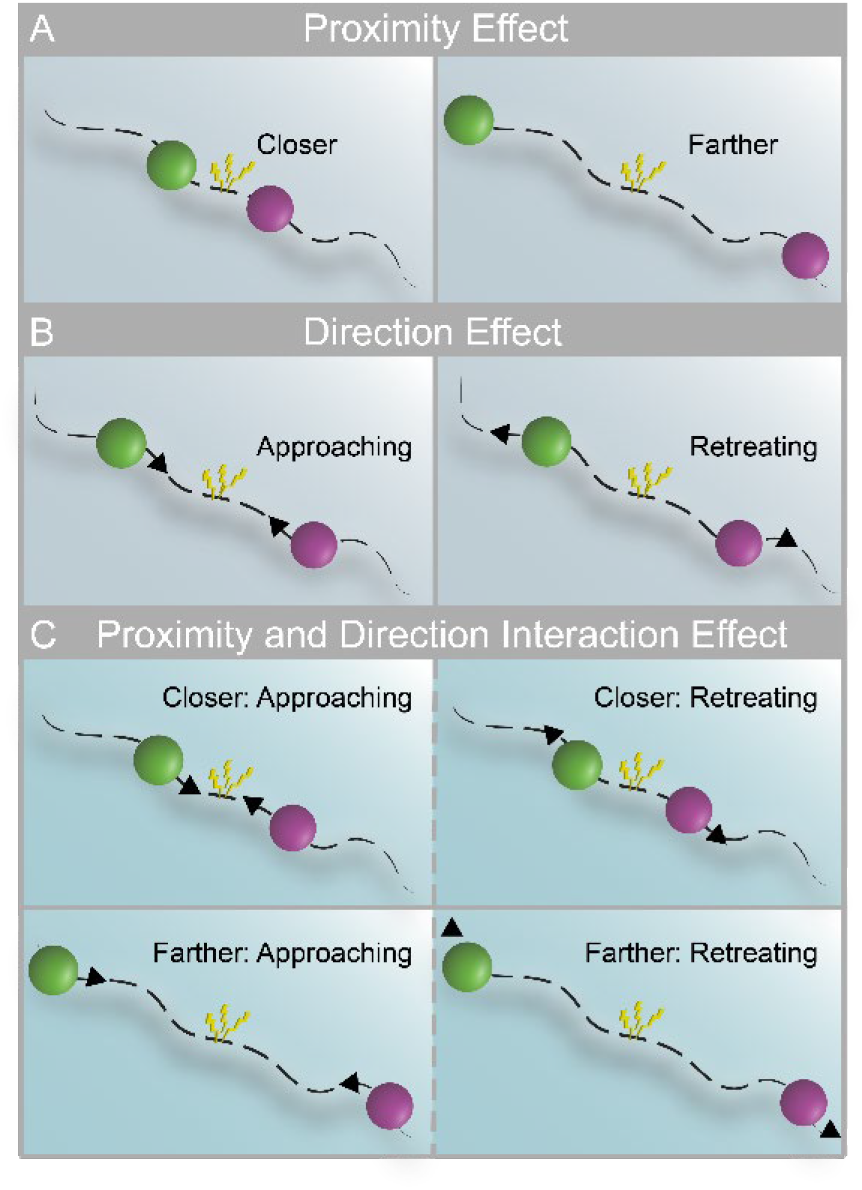
Threat-related factors and their interaction. (A) Closer and farther threat, where threat is represented by an aversive shock when circles touched. (B) Direction of threat: approach vs. retreat. (C) Threat level may depend on both proximity (closer and farther) and direction (left panels indicate approach; right panels indicate retreat).

Thus far, studies have considered the effects of threat proximity and direction independently. Hence, it is currently unknown how such factors potentially interact in the brain during aversive anticipation (Figure 1C). This is an important gap in our knowledge base because behavioral findings have extensively documented interactions between threat-related factors, which have produced several influential theoretical accounts (for excellent discussion, see Mobbs et al., 2015). Furthermore, it is not only important to investigate how multiple threat-related factors interact but to understand how the brain tracks them continuously. In particular, do signal fluctuations in brain regions track threat-related factors dynamically? If so, to what factor(s) and factor combinations are they sensitive?

To address these questions, we devised a paradigm in which threat was dynamically modulated during fMRI scanning. Two circles moved on the screen, sometimes moving closer and sometimes moving apart, and at varying speeds (Figure 2). Participants were instructed to pay attention to the circles on the screen and were explicitly informed that, if they touched, the participants would receive an unpleasant shock. As a measure of threat-evoked physiological arousal, skin conductance responses were recorded during scanning. Our paradigm allowed us to investigate the role played by the interaction between proximity (nearer vs. farther circles), direction (approach vs. retreat), and speed (faster vs. slower) in determining brain responses during anticipatory threat processing. Importantly, the impact of the factors “proximity” and “speed” were assessed parametrically (i.e., continuously) as they varied dynamically. Therefore, the paradigm allowed us to test how multiple threat-related factors *dynamically* influence signals fluctuations across brain regions. Specifically, do they provide independent contributions or do they interact in regions important for threat processing, such as the anterior insula, amygdala, PAG, and BST? Intuitively, probing interactions allowed us to evaluate the extent to which the influence of one factor on threat anticipation depended on the values of other factor(s). For instance, in terms of a two-way interaction, we anticipated that the influence of direction (i.e., approaching vs. retreating threat) would depend on proximity (i.e., whether the threat was near vs. far; Figure 1C). In terms of three-way interactions, we sought to evaluate if the interaction between the continuously manipulated factors of proximity and speed depended on direction.

**Figure 2.**
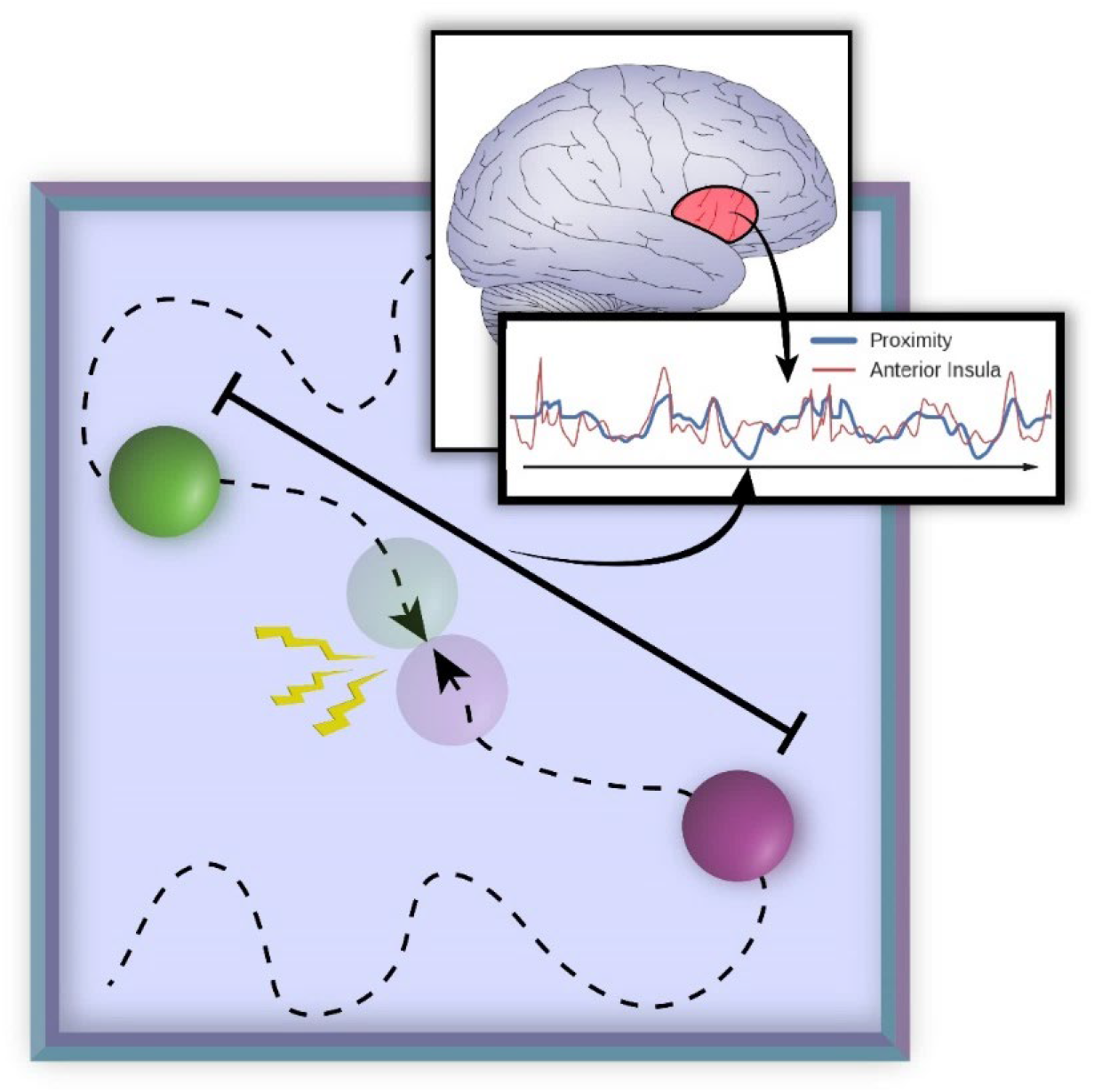
Experimental paradigm. Two circles moved randomly on the screen and a shock was administered to the participant if they touched. The inset represents threat proximity (the distance between the two circles), which varied continuously. A central goal of the study was to determine the extent to which signal fluctuations in brain regions (such as the anterior insula) followed threat-related factors (including proximity) and their interactions.

## Materials and Methods

### Participants

Eighty-five participants (41 females, ages 18-40 years; average: 22.62, STD: 4.85) with normal or corrected-to-normal vision and no reported neurological or psychiatric disease were recruited from the University of Maryland community (of the original sample of 93, data from 7 subjects were discarded due to technical issues during data transfer [specifically, field maps were lost] and 1 other subject was removed because of poor structural-functional alignment). The project was approved by the University of Maryland College Park Institutional Review Board and all participants provided written informed consent before participation. The data analyzed here were investigated in an entirely separate fashion at the level of networks and published previously (Najafi et al., 2017). The sample size was not based on an explicit statistical power analysis. At the outset, we sought to collect around 90 participants to allow investigation of the data in terms of separate “exploratory” and “test” sets in the network study (Najafi et al., 2017). For the investigation of activation (present paper), our intention was to employ the available data in a single type of analysis.

### Anxiety questionnaires

Participants completed the trait portion of the Spielberger State-Trait Anxiety Inventory (STAI; Spielberger et al., 1970) before scanning (average: 17.23 days, STD: 15.90), and then completed the state portion of the STAI immediately before the scanning session.

### Procedure and Stimuli

Two circles with different colors moved around on the screen randomly. When they collided with each other, an unpleasant mild electric shock was delivered. Overall, proximity, direction of movement, and relative speed of the circles were used to influence perceived threat. The position of each circle (on the plane), ***x***_*t*_, was defined based on its previous position, ***x***_*t*−1_, plus a random displacement, ∆***x***_*t*_:

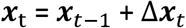

The magnitude and direction of the displacement was calculated by combining a normal random distribution with a momentum term to ensure motion smoothness, while at the same time remaining (relatively) unpredictable to the participants. Specifically, the displacement was updated every 50 ms as follows:

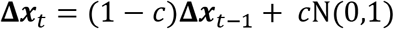

where *cc* = 0.2 and N(0,1) indicates the normal distribution with zero mean and standard deviation of 1.

Visual stimuli were presented using PsychoPy (http://www.psychopy.org/) and viewed on a projection screen via a mirror mounted to the scanner’s head coil. Each participant viewed the same sequence of circle movements. The total experiment included 6 runs (457 seconds each), each of which had 6 blocks (3/85 participants had only 5 runs). In each block, the circles appeared on the screen and moved around for 60 seconds; blocks were separated by a 15-second off period during which the screen remained blank. Each run ended with a 7-second blank screen.

To ensure that the effects of threat proximity and direction were uncorrelated, half of the blocks in each run were temporally reversed versions of the other blocks in that run. Temporally reversing the stimulus trajectories guarantees that proximity and direction are uncorrelated because reversing time changes the sign of the direction (that is, approach becomes retreat). To optimize the experimental design, 10,000 candidate stimuli trajectories and block orders were generated. We then selected six runs which minimized collinearity between all predictors of interest (see below), measured as the sum of respective variance inflation factors (Neter et al., 1996).

In each run the circles collided 8 times within 4 out of 6 blocks (1-3 times in a block); in the remaining 2 blocks there were no collisions. Each collision resulted in the delivery of an electric shock. The 500-ms electric shock (comprised of a series of current pulses at 50 Hz) was delivered by an electric stimulator (Model number E13-22 from Coulbourn Instruments, PA, USA) to the fourth and fifth fingers of the non-dominant left hand via MRI-compatible electrodes. To calibrate the intensity of the shock, each participant was asked to choose his/her own stimulation level immediately prior to functional imaging, such that the stimulus would be “highly unpleasant but not painful.” After each run, participants were asked about the unpleasantness of the stimulus in order to re-calibrate shock strength, if needed. Skin conductance response (SCR) data were collected using the MP-150 system (BIOPAC Systems, Inc., CA, USA) at a sampling rate of 250 Hz by using MRI compatible electrodes attached to the index and middle fingers of the non-dominant left hand. Due to technical problems and/or experimenter errors during data collection, SCR data was not available in 2 participants, and 6 participants had only 5 runs of the SCR data; 1 participant who had only 3 runs of data was excluded from the analysis of SCR data.

### MRI data acquisition

Functional and structural MRI data were acquired using a 3T Siemens TRIO scanner with a 32-channel head coil. First, a high-resolution T2-weighted anatomical scan using Siemens’s SPACE sequence (0.8 mm isotropic) was collected. Subsequently, we collected 457 functional EPI volumes in each run using a multiband scanning sequence (Feinberg et al., 2010) with TR = 1.0 sec, TE = 39 ms, FOV = 210 mm, and multiband factor = 6. Each volume contained 66 non-overlapping oblique slices oriented 30° clockwise relative to the AC-PC axis (2.2 mm isotropic). A high-resolution T1-weighted MPRAGE anatomical scan (0.8 mm isotropic) was collected. Additionally, in each session, double-echo field maps (TE1 = 4.92 ms, TE2 = 7.38 ms) were acquired with acquisition parameters matched to the functional data.

### Functional MRI preprocessing

To preprocess the functional and anatomical MRI data, a combination of packages and in-house scripts were used. The first three volumes of each functional run were discarded to account for equilibration effects. Slice-timing correction (with Analysis of Functional Neuroimages’ (AFNI; Cox, 1996) 3dTshift) used Fourier interpolation to align the onset times of every slice in a volume to the first acquisition slice, and then a six-parameter rigid body transformation (with AFNI’s 3dvolreg) corrected head motion within and between runs by spatially registering each volume to the first volume.

In this study, we strived to improve functional-anatomical co-registration given the small size of some of the structures of interest. Skull stripping determines which voxels are to be considered part of the brain and, although conceptually simple, plays a very important role in successful subsequent co-registration and normalization steps. Currently, available packages perform sub-optimally in specific cases, and mistakes in the brain-to-skull segmentation can be easily identified. Accordingly, to skull strip the T1 high-resolution anatomical image (which was rotated to match the oblique plane of the functional data with AFNI’s 3dWarp), we employed six different packages [ANTs (Avants et al., 2009; http://stnava.github.io/ANTs/), AFNI (Cox, 1996; http://afni.nimh.nih.gov/), ROBEX (Iglesias et al., 2011; https://www.nitrc.org/projects/robex), FSL (Smith et al., 2004; http://fsl.fmrib.ox.ac.uk/fsl/fslwiki/), SPM (http://www.fil.ion.ucl.ac.uk/spm/), and BrainSuite (Shattuck and Leahy, 2002; http://brainsuite.org/)] and employed a “voting scheme” as follows: based on T1 data, a voxel was considered to be part of the brain if 4/6 packages estimated it to be a brain voxel; otherwise the voxel was not considered to be brain tissue (for 6 subjects whose T1 data were lost due to issues during data transfer, the T2 image was used instead and only the ANTs package was used for skull-stripping).

Subsequently, FSL was used to process field map images and create a phase-distortion map for each participant (by using bet and fsl_prepare_fieldmap). FSL’s epi_reg was then used to apply boundary-based co-registration to align the unwarped mean volume registered EPI image with the skull-stripped anatomical image (T1 or T2), along with simultaneous EPI distortion-correction (Greve and Fischl, 2009).

Next, ANTS was used to learn a nonlinear transformation that mapped the skull-stripped anatomical image (T1 or T2) to the skull-stripped MNI152 template (interpolated to 1-mm isotropic voxels). Finally, ANTS combined the nonlinear transformations from co-registration/unwarping (from mapping mean functional EPI image to the anatomical T1 or T2) and normalization (from mapping T1 or T2 to the MNI template) into a single transformation that was applied to map volume-registered functional volumes to standard space (interpolated to 2-mm isotropic voxels). In this process, ANTS also utilized the field maps to simultaneously minimize EPI distortion. The resulting spatially normalized functional data were blurred using a 4mm full-width half-maximum (FWHM) Gaussian filter. Spatial smoothing was restricted to grey-matter mask voxels. Finally, intensity of each voxel was normalized to a mean of 100 (separately for each run).

### Voxelwise analysis

Each participant’s preprocessed functional MRI data were analyzed using multiple linear regression with AFNI (restricted to gray-matter voxels) using the 3dDeconvolve program (https://afni.nimh.nih.gov/afni/doc/manual/3dDeconvolve.pdf). Time series data were analyzed according to the following model (additional nuisance variables are described below):

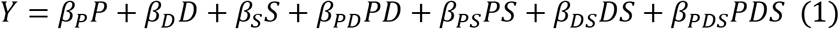

where *P* indicates proximity, *D* represents direction, and *S* represents speed. Variables were determined based on circle positions on the screen. Proximity was defined as the Euclidean distance between the two circles; direction indicated approach vs. retreat; speed was the discrete temporal difference of proximity. The products *PD*, *PS*, and *PDS* represent the interactions terms; the individual terms *P*, *D*, and *S* were mean centered prior to multiplication to reduce potential collinearity. The resulting regressors exhibited pairwise correlations that were relatively small (the largest was .41) and all variance inflation factors were less than 1.3, indicating that model estimation was unproblematic (Mumford et al., 2015).

In addition to the variables above, we included regressors for visual motion (velocity tangential to the difference vector of the combined circle-to-circle stimulus), sustained block event (60-sec duration), and block-onset and block-offset events (1-second duration) to account for transient responses at block onset/offset. All regressors were convolved with a standard hemodynamic response based on the gamma-variate model (Cohen, 1997). Note that interaction regressors were multiplied prior to convolution; also, as stimulus-related display information was updated every 50 ms (20 Hz), convolution with the hemodynamic response was performed prior to decimating the convolved signal to the fMRI sample rate (1 Hz). To simplify plotting, decimated regressors were scaled by their corresponding root mean square value (thus, multiplicative interactions terms were on the same scale as simple effects). Other regressors included in the model included 6 motion parameters (3 linear displacements and 3 angular rotations), and their discrete temporal derivatives. To further control for head motion-related artifacts in the data (Siegel et al., 2014), we excluded volumes (on average 0.4%) with a frame-to-frame displacement of more than 1 mm. To model baseline and drifts of the MRI signal, regressors corresponding to polynomial terms up to 4th order were included (for each run separately). Finally, to minimize effects due to the physical shock event, data points in a 15-sec window after shock delivery were discarded from the analysis. It should be pointed out that to partly account for the fact that the circles were most proximal just prior to shock events, the design included time periods when circles were very close but did not touch eventually.

### Group analysis

Whole-brain voxelwise random-effects analyses were conducted using response estimates from individual-level analyses (restricted to gray-matter voxels) in AFNI. To probe the effects of the regressors of interest, we ran separate one-sample *t*-tests against zero using the AFNI’s 3dttest++ program.

The alpha-level for voxelwise statistical analysis was determined by simulations using the 3dClustSim program (restricted to gray-matter voxels). For these simulations, the smoothness of the data was estimated using 3dFWHMx program (restricted to gray-matter voxels) based on the residual time series from the individual-level voxelwise analysis. Taking into account the recent report of increased false-positive rates linked to the assumption of Gaussian spatial autocorrelation in fMRI data (Eklund et al., 2016), we used the -acf (i.e., auto-correlation function) option recently added to the 3dFWHMx and 3dClustSim tools, which models spatial fMRI noise as a mixture of Gaussian plus mono-exponential distributions. This improvement was shown to control false positive rates around the desired alpha level, especially with relatively stringent voxel-level uncorrected p-values such as 0.001 (Cox et al., 2017). Based on a voxel-level uncorrected p-value of 0.001, simulations indicated a minimum cluster extent of 13 voxels (2.0 × 2.0 × 2.0 mm) for a cluster-level corrected alpha of 0.05.

### BST ROI analysis

The BST is a basal forebrain region and has been frequently implicated in threat-related processing (Davis et al., 2009; Fox et al., 2015) along with other regions such as the amygdala and anterior insula (Pessoa, 2016). Because the BST is a small region, analysis based on spatially smoothed data would be susceptible to signals from surrounding structures. To reduce this possibility, we conducted an additional BST ROI analysis using spatially unsmoothed data. Bilateral BST ROIs were defined anatomically according to the probabilistic mask of the BST (at 25% threshold) recently reported by Blackford and colleagues (Theiss et al., 2017). For this analysis, no spatial smoothing was applied. In each participant, for each ROI, a representative time series was created by averaging the unsmoothed time series from all the gray-matter voxels within the anatomically defined ROI (left: 9 voxels; right: 8 voxels). Then, as in the individual-level voxelwise analysis, multiple linear regression analysis was run using the 3dDeconvolve program to estimate condition-specific responses. At the group level, as in the voxelwise analysis, we ran separate one-sample *t*-test’s against zero using the corresponding regression coefficients from the individual-level analysis.

### Skin conductance response (SCR) analysis

Each participant’s SCR data were initially smoothed with a median-filter over 50 samples (200 ms) to reduce scanner-induced noise. In each run, the first 3 seconds of data were discarded (corresponding to first 3 volumes excluded in the fMRI analysis) and the remaining data were resampled by decimating the 250 Hz sample rate to the sample rate of fMRI data (1 Hz) and subsequently *Z*-scored. The pre-processed SCR data were then analyzed using multiple linear regression using the 3dDeconvolve program in AFNI (for related approaches see Bach et al. (2009) and Engelmann et al. (2015)). We employed the same regression model as the one used for fMRI data (see equation 1). In addition, we included regressors for visual motion (velocity tangential to the difference vector of the combined circle-to-circle stimulus), sustained block event (60-sec duration), and block-onset and block-offset events (1-second duration) to account for transient responses at block onset/offset. All regressors were convolved with a canonical skin conductance response model based on the sigmoid-exponential function (Lim et al., 1997; Figure 3). Additionally, constant and linear terms were included (for each run separately) to model baseline and drifts of the SCR. To minimize effects due to the physical shock event, data points in a 15-sec window after shock delivery were discarded from the analysis. At the group level, to probe the effects of the regressors of interest, we ran separate one-sample *t*-tests against zero using the corresponding regression coefficients from the individual-level analysis.

**Figure 3.**
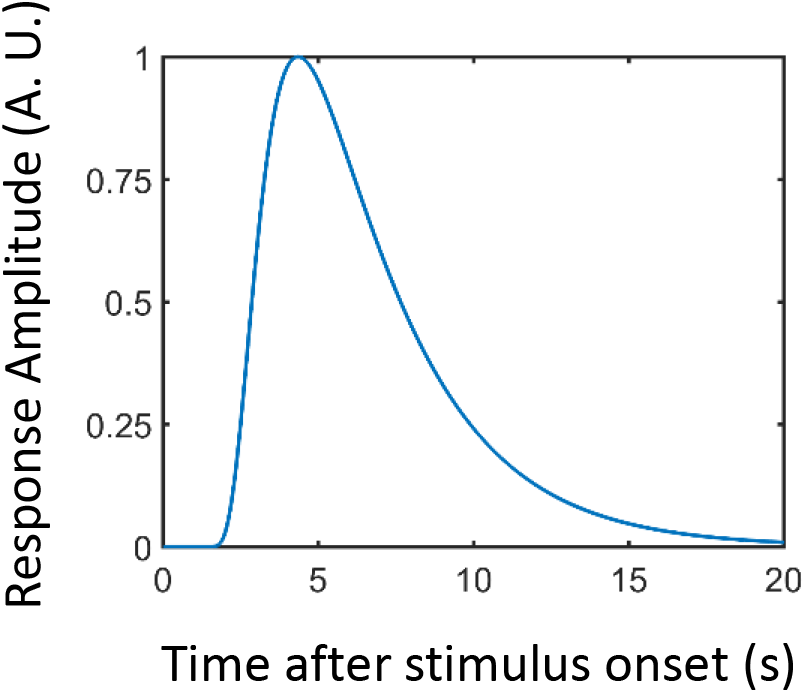
Skin conductance response (SCR) model based on the sigmoid-exponential function (Lim et al., 1997). A.U.: arbitrary units.

### Relationship between SCR and brain activity

To probe the relationship between brain activity and physiological arousal, we focused on the right anterior insula and the right amygdala clusters that exhibited a proximity by direction interaction (see Results). For each cluster, an interaction index was created by averaging the corresponding regression coefficients (*β_PD_* in equation 1) from all the voxels within the cluster (after cluster-level thresholding). Then, for each cluster, we ran a robust correlation (Rousselet and Pernet, 2012; Wilcox, 2012) across participants. For each participant, we considered the average fMRI interaction regression coefficient and the corresponding interaction term in the SCR data (specifically, the coefficient *β_PD_* obtained from the SCR regression analysis).

### Relationship between threat anticipation and physical shock responses

In an exploratory analysis, we probed the relationship between activity related to threat anticipation and responses to physical shock itself. For the anticipatory activity, we considered the proximity by direction interaction and focused on the right anterior insula and right amygdala clusters which exhibited this interaction (see Results). To estimate responses to physical shocks, we ran a separate multiple regression analysis with all the regressors as in the original model along with an additional regressor that modeled physical shock events (500 ms). As noted above, these events were discarded in the main analyses to minimize potential contributions from actual electrical stimulation. Then, for each cluster, we ran a robust correlation (Rousselet and Pernet, 2012; Wilcox, 2012) across participants. For each participant, we considered the average regression coefficient corresponding to the proximity by direction interaction (from the original model so as to estimate it with minimal contamination from shocks) and regression coefficient corresponding to physical shock events.

### Plotting parametric effects as a function of proximity

Equation 1 allowed us to estimate the contributions of the seven main regressors to fMRI responses. Because of the parametric nature of the design, to illustrate responses in a more intuitive manner, we estimated responses separately for approach and retreat for a range of proximity values (Figure 8). To do so, the value of *z*-scored proximity was varied (in the range of [−2, 1.5] and at the mean speed value), and the estimated regression coefficients were used to estimate the response at each value of proximity.

To provide an indication of variability of the fit across participants, we adopted the following approach. In the case of the proximity by direction interaction (Figures 8 and 11A), at each level of proximity, we calculated the difference between the estimated response for the approach and retreat conditions. We then calculated the standard error of the approach-minus-retreat difference across participants (at each value of proximity). We display the 95% confidence bands at each proximity value (note that because the intervals were based on differences between approach and retreat conditions, the same band widths are employed for approach and retreat). An analogous procedure was employed for the proximity by direction interaction of SCRs (Figure 4). The BST exhibited a proximity effect but no interaction. Therefore, in Figure 9 we computed error bands separately for approach and retreat based on the variability of estimated responses across participants as a function of proximity.

**Figure 4.**
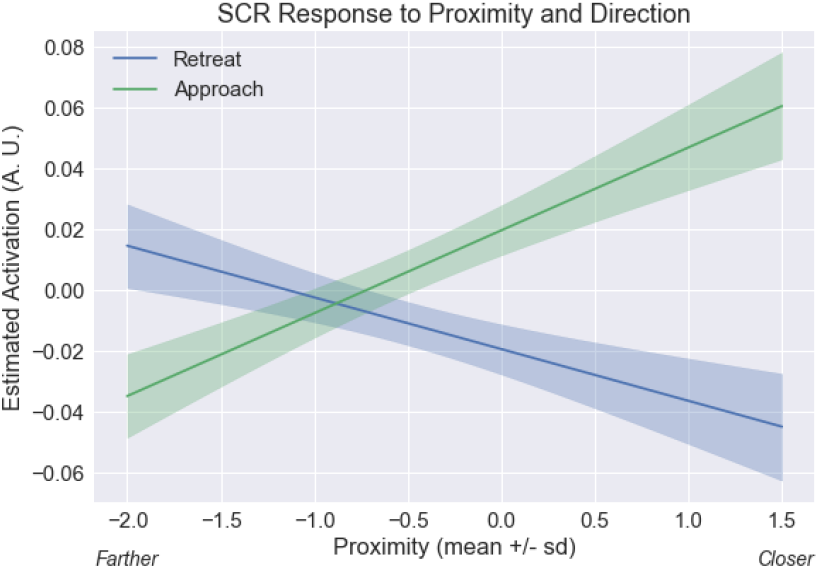
Skin conductance response (SCR) proximity by direction interaction. Estimated responses for a range of proximity values. To display estimated responses, we varied proximity and estimated the response based on the linear model for SCR (analogous to the model of equation 1). The approach vs. retreat difference was greater when circles were near compared to far. The confidence bands were obtained by considering within-subject differences (approach minus retreat); see Methods. A.U.: arbitrary units.

### Statistical approach and p values

The null hypothesis significance testing framework has come under increased scrutiny in recent years. In particular, the hard threshold of .05 has come under attack, with reasonable researchers calling for both stricter thresholds (Benjamin et al., 2017) or, conversely, for p values to be abandoned (McShane et al., 2017). However, like McShane and colleagues, we do not consider a binary threshold to be satisfactory, and believe that p values should be treated continuously. Accordingly, in select cases, we show p values and discuss findings that do not survive correction for multiple comparisons; in the context of Table 9, we discuss the general results of the BST given its important role in threat-related processing.

## Results

Our paradigm allowed us to investigate the role played by threat proximity, direction, and speed, and their interactions, on SCRs and fMRI responses. Intuitively, interactions evaluated the extent to which factor combinations were relevant in explaining the data. For instance, the contrast of approach vs. retreat (direction) was anticipated to depend on proximity (Figure 1C). Moreover, as proximity and speed varied continuously, their roles and their interactions were assessed parametrically.

Our design did not include a standard control condition (for example, circles colliding but no shock administered), as often is the case in fMRI studies. Note, however, that our main goal was not to investigate the shock event itself but potential threat. Thus, approach and retreat can be viewed as paired conditions insofar as processes related to tracking the movement of the circles are concerned, for example. Furthermore, as stated in the preceding paragraph, an important focus of the research was to assess whether or not brain regions were sensitive to variable interactions, an approach that further helped reduce the contributions of non-threat related processing (see also Discussion).

### Skin Conductance Responses

Analysis of SCR data revealed that all three main variables had robust effects on responses (Table 1). In addition, we detected an interaction of proximity by direction; in this case responses to approach vs. retreat were sensitive to threat distance, such that the effect was larger when near vs. far. To visualize this result, Figure 4 shows estimated SCRs for approach and retreat for a range of proximity values (because the circles moved continuously on the screen, Figure 4 employed an approach similar to that of Figure 8 for plotting; see Methods). Finally, a three-way interaction between proximity, direction, and speed also survived correction for multiple comparisons.

**Table 1.** Skin conductance response results.

| **Regressor** | **t(81)** | **p-value** |
| --- | --- | --- |
| Proximity | 4.57 | 0.0000 |
| Direction | 9.37 | 0.0000 |
| Speed | -4.20 | 0.0001 |
| DirectionXSpeed | -0.92 | 0.3602 |
| ProximityXDirection | 10.99 | 0.0000 |
| ProximityXSpeed | -2.43 | 0.0175 |
| ProximityXDirectionXSpeed | -2.78 | 0.0067 |

**Bonferroni correction for multiple comparisons: $0.05/7 = 0.0071$**

### fMRI voxelwise analysis

Figures 5-6 (Tables 2-3) show the effects of proximity and direction (Table 4 shows the effect of speed). The main focus of this study was to investigate interactions between threat-related factors. Figure 7 (Table 5) shows interactions between proximity and direction; positive voxels (red) show effects when the contrast of approach vs. retreat was greater during closer vs. farther circles, and blue voxels indicate the opposite. Figure 8 shows estimated responses for approach and retreat for a range of proximity values, which aids in visualizing the parametric effects of proximity on the signals in the two regions (see Methods). For the right anterior insula (Figure 8A), when the circles were closer to each other, a larger approach vs. retreat differential response was observed compared to when the circles were farther from each other. Responses for the right amygdala (Figure 8B) exhibited the opposite pattern as responses were larger for retreat compared to approach, and the contrast was enhanced when circles were closer compared to farther. Tables 6-7 show two-way interactions between direction and speed and between proximity and speed. Table 8 shows the three-way interaction of proximity, direction and speed.

**Figure 5.**
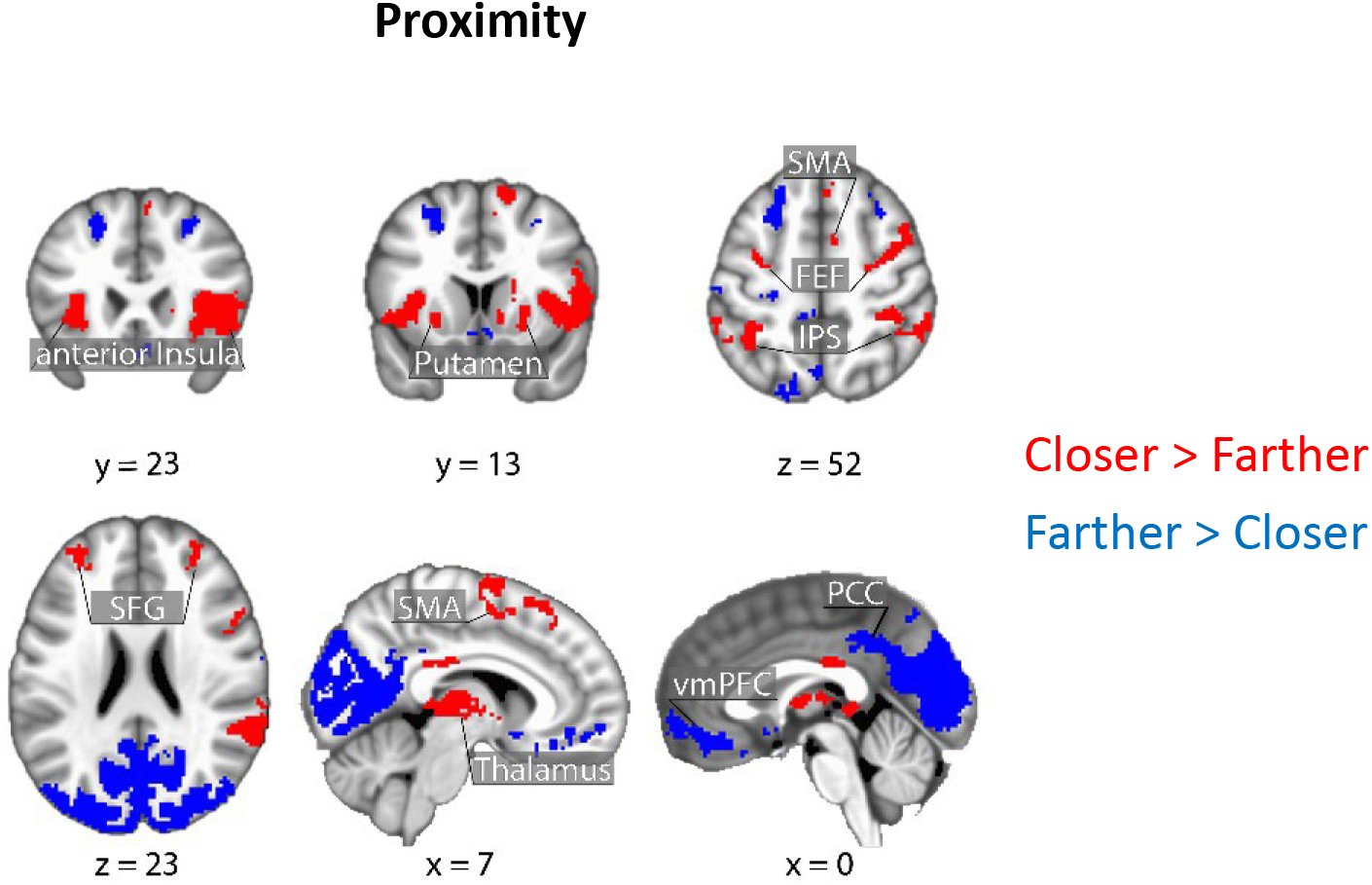
Brain responses as a function of threat proximity. Clusters in red show regions with stronger responses for closer vs. farther; clusters in blue show the reverse. Clusters were thresholded at a whole-brain corrected alpha of .05. SMA: supplementary motor area; FEF: frontal eye field; IPS: intraparietal sulcus; SFG: superior frontal gyrus; PCC: posterior cingulate cortex; vmPFC: ventro-medial prefrontal cortex.

**Figure 6.**
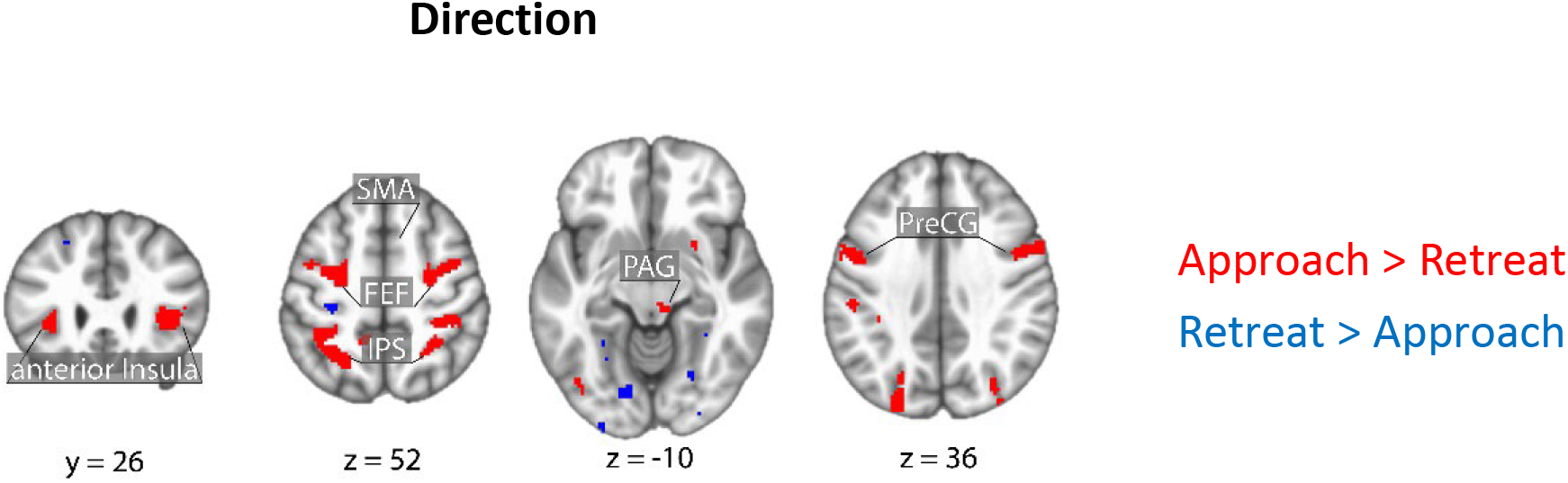
Brain responses as a function of direction (approach vs. retreat). Clusters in red show regions with stronger responses for approach vs. retreat; clusters in blue show the reverse. Clusters were thresholded at a whole-brain corrected alpha of .05. PAG: periaqueductal gray; SMA: supplementary motor area; FEF: frontal eye field; IPS: intraparietal sulcus; PreCG: precentral gyrus.

**Figure 7.**
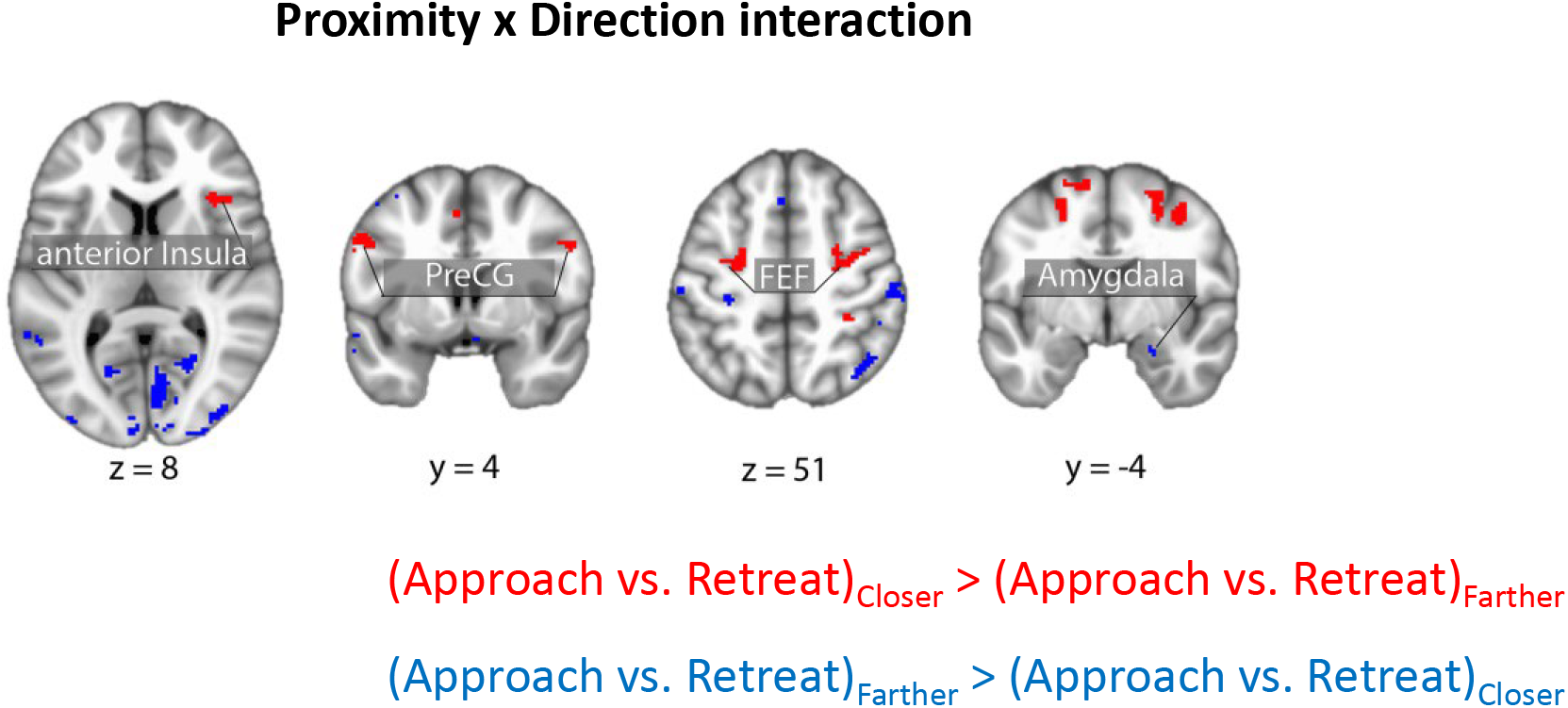
Brain responses exhibiting a proximity by direction (approach vs. retreat) interaction in areas of interest. Clusters in red show regions with approach vs. retreat responses greater when closer vs. farther; clusters in blue show the reserve pattern. Clusters were thresholded at a whole-brain corrected alpha of .05. FEF: frontal eye field; PreCG: precentral gyrus.

**Figure 8.**
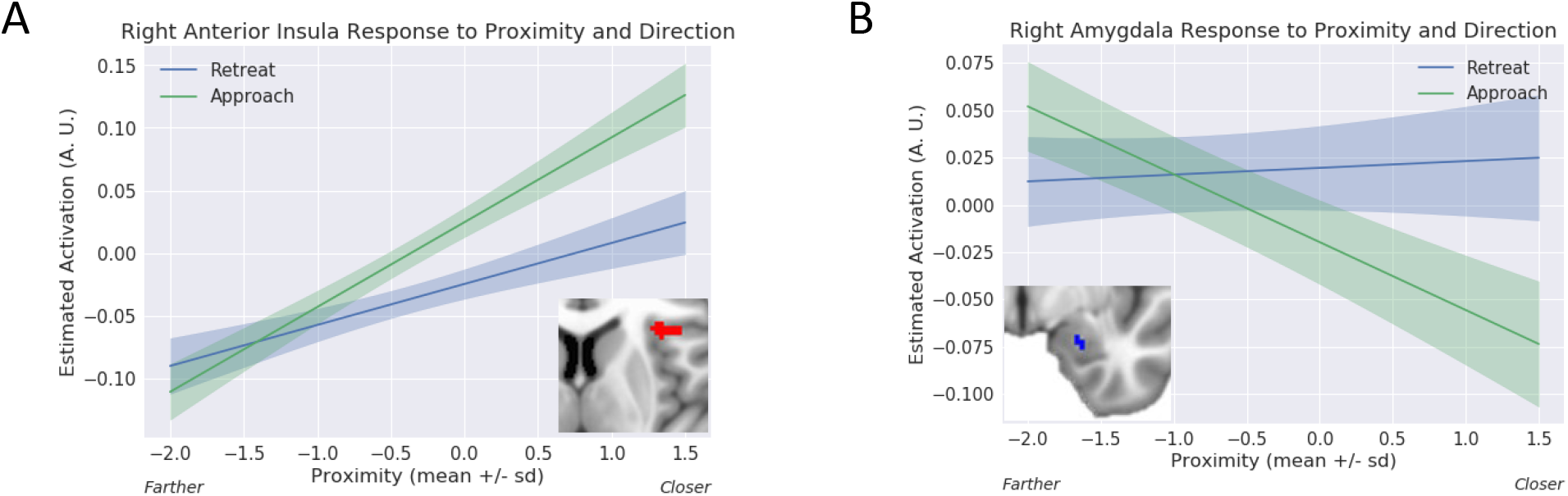
Proximity by direction (approach vs. retreat) interaction. Estimated responses for a range of proximity values. (A) For the right anterior insula, activity increased as a function of proximity for both approach and retreat, but more steeply for the former. (B) For the right amygdala, activity decreased as a function of proximity during approach, but changed little during retreat. The confidence bands were obtained by considering within-subject differences (approach minus retreat); see Methods. A.U.: arbitrary units.

**Table 2.** Clusters that exhibited the effect of proximity in voxelwise analysis at whole-brain cluster-level corrected alpha of 0.05 (peak MNI coordinates, t(84) values, and cluster size [k] refers to number of 2.0 × 2.0 × 2.0 mm^3^ voxels). Peak coordinates are presented for completeness and potential meta-analysis; with cluster-based thresholding, it is not possible to conclude that all the reported peaks were activated [see Woo et al., 2014]

| k | x | y | z | t | Cluster |
| --- | --- | --- | --- | --- | --- |
| 12470 | -14 | -88 | 28 | -13.47 | Occipital cortex/Cuneus/Posterior Cingulate cortex |
| 1690 | 36 | 22 | 8 | 7.93 | right anterior/mid-Insula |
| 1489 | -34 | -92 | -6 | 8.84 | left inferior/middle Occipital gyrus |
| 1453 | 28 | -90 | 4 | 8.93 | right inferior/middle Occipital gyrus |
| 1188 | 64 | -38 | 30 | 7.83 | right Supramarginal/Postcentral gyrus |
| 1088 | 14 | 10 | 64 | 6.40 | right Superior Frontal gyrus |
| 995 | -16 | -76 | -34 | 7.84 | left Cerebellum |
| 869 | -60 | -46 | 42 | 6.58 | left Supramarginal gyrus |
| 796 | -32 | 22 | 6 | 7.28 | left Anterior Insula |
| 576 | -2 | 46 | -10 | -5.91 | ventro-medial Prefrontal Cortex |
| 526 | 8 | -18 | 6 | 7.49 | right/left Thalamus |
| 336 | -22 | 26 | 46 | -5.23 | left Superior Frontal gyrus |
| 333 | -12 | -72 | -44 | 5.89 | left Cerebellum |
| 209 | 44 | -58 | -30 | 5.88 | right Cerebellum |
| 138 | 22 | 32 | 48 | -4.55 | right Superior Frontal gyrus |
| 125 | -4 | -20 | 30 | 5.27 | right/left posterior Cingulate cortex |
| 118 | -34 | 48 | 28 | 4.96 | left Middle Frontal gyrus |
| 117 | -62 | -6 | -14 | -5.84 | left Middle Temporal gyrus |
| 117 | 18 | 6 | 18 | 5.41 | right dorso-lateral Caudate |
| 88 | 26 | 42 | 22 | 4.47 | right Middle Frontal gyrus |
| 83 | -26 | 6 | -10 | 5.18 | left Putamen |
| 79 | -12 | -54 | 66 | -5.71 | left Precuneus/Superior parietal lobule |
| 72 | 4 | 32 | 48 | 4.88 | medial Superior Frontal gyrus |
| 70 | 58 | -30 | -6 | 4.36 | right Superior/Middle Temporal gyrus |
| 68 | 56 | -4 | -18 | -4.98 | right Middle Temporal gyrus |
| 66 | -26 | -24 | 54 | -4.83 | left Precentral gyrus |
| 65 | -34 | -6 | 50 | 4.57 | left Middle frontal gyrus |
| 61 | 42 | -12 | 48 | -4.85 | right Precentral gyrus |
| 55 | -42 | -24 | 18 | -4.11 | left Posterior Insula |
| 42 | 32 | 40 | -10 | -5.02 | right lateral Orbitofrontal cortex |
| 40 | -12 | -2 | 66 | 4.73 | left Superior Frontal gyrus |
| 36 | -58 | -22 | 48 | -4.09 | left Postcentral gyrus |
| 34 | 36 | -70 | -10 | 5.04 | right Inferior Temporal gyrus |
| 34 | 4 | 12 | -10 | -5.17 | Subcallosal area |
| 34 | -4 | -68 | 50 | -4.38 | Precuneus |
| 31 | 20 | -14 | -24 | -5.47 | right Hippocampus/Amygdala |
| 25 | -22 | -8 | -22 | -4.36 | left Hippocampus/Amygdala |
| 24 | -4 | -24 | 54 | -4.52 | left Paracentral lobule |
| 20 | 38 | -12 | -8 | 4.36 | right Planum Polare |
| 20 | 54 | -26 | 10 | -4.20 | right Transverse Temporal gyrus |
| 17 | 12 | 12 | -2 | 5.02 | right ventral Caudate |
| 17 | -52 | -28 | 10 | -3.91 | left Transverse Temporal gyrus |
| 17 | -16 | 64 | 14 | -4.22 | right Superior Frontopolar gyrus |
| 17 | 66 | -4 | 18 | -4.78 | right Precentral gyrus |
| 17 | 14 | -54 | 68 | -4.57 | right Precuneus/Superior parietal lobule |
| 16 | -26 | 62 | -12 | 4.42 | left frontomarginal gyrus |
| 15 | 12 | -74 | -40 | 5.49 | right Cerebellum |
| 14 | -56 | -50 | -10 | -4.11 | left Superior/Middle Temporal gyrus |
| 13 | -18 | -34 | 72 | -4.34 | left Postcentral gyrus |

**Table 3.**
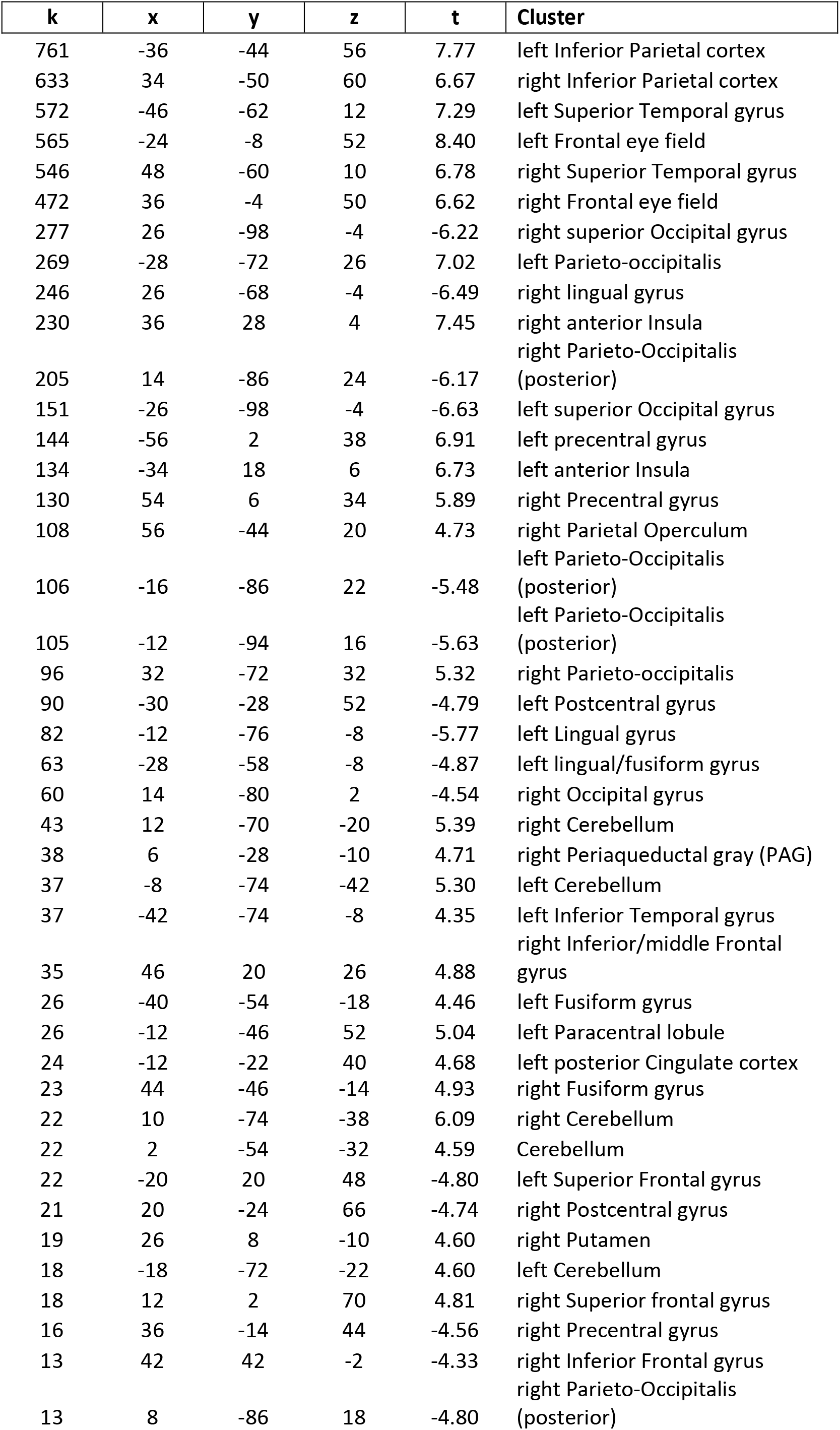
Clusters that exhibited the effect of direction in voxelwise analysis at whole-brain cluster-level corrected alpha of 0.05 (peak MNI coordinates, t(84) values, and cluster size [k] refers to number of 2.0 × 2.0 × 2.0 mm^3^ voxels). Peak coordinates are presented for completeness and potential meta-analysis; with cluster-based thresholding, it is not possible to conclude that all the reported peaks were activated [see Woo et al., 2014]

**Table 4.**
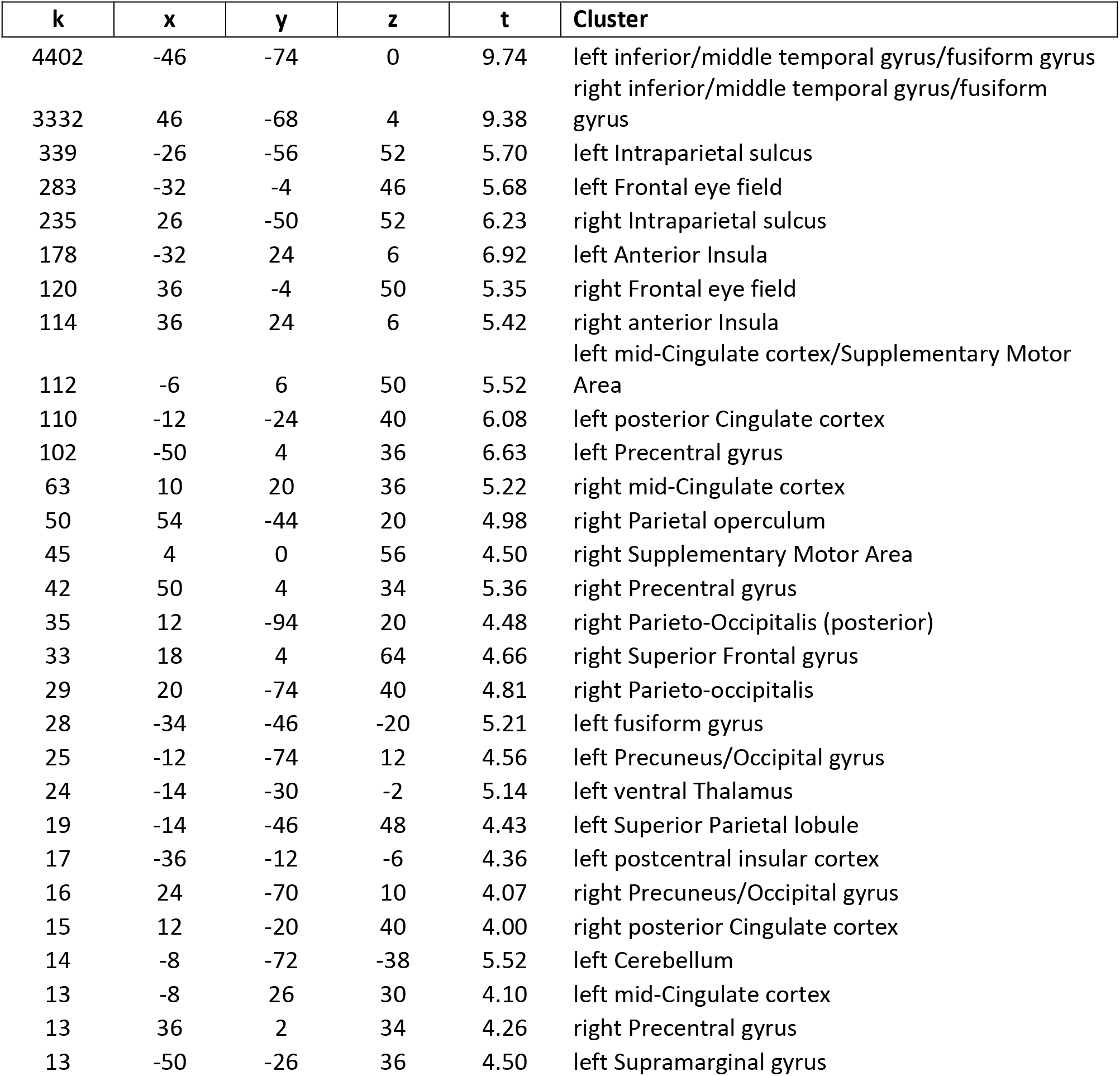
Clusters that exhibited the effect of speed in voxelwise analysis at whole-brain cluster-level corrected alpha of 0.05 (peak MNI coordinates, t(84) values, and cluster size [k] refers to number of 2.0 × 2.0 × 2.0 mm^3^ voxels). Peak coordinates are presented for completeness and potential meta-analysis; with cluster-based thresholding, it is not possible to conclude that all the reported peaks were activated [see Woo et al., 2014]

**Table 5.** Clusters that exhibited the proximity x direction interaction in voxelwise analysis at whole-brain cluster-level corrected alpha of 0.05 (peak MNI coordinates, t(84) values, and cluster size [k] refers to number of 2.0 × 2.0 × 2.0 mm^3^ voxels). Peak coordinates are presented for completeness and potential meta-analysis; with cluster-based thresholding, it is not possible to conclude that all the reported peaks were activated [see Woo et al., 2014]

| **k** | **x** | **y** | **z** | **t** | **Cluster** |
| --- | --- | --- | --- | --- | --- |
| 528 | 10 | -74 | -2 | -9.42 | right Occipital cortex |
| 398 | -10 | -96 | 12 | -7.09 | left Occipital gyrus |
| 358 | 28 | -96 | 2 | -5.98 | right Occipital gyrus |
| 263 | -20 | -12 | 60 | 6.26 | left Frontal eye field |
| 243 | 16 | -86 | 26 | -7.16 | right Occipital gyrus |
| 215 | 26 | -2 | 58 | 5.55 | right Frontal eye field |
| 159 | -54 | -32 | -2 | -5.16 | left Superior Temporal gyrus |
| 148 | -16 | -86 | 22 | -5.94 | left Occipital gyrus |
| 138 | -28 | -98 | 0 | -5.24 | left inferior Occipital gyrus |
| 109 | -54 | 4 | 38 | 6.72 | left Precentral gyrus |
| 109 | 50 | -32 | 58 | -4.88 | right Postcentral gyrus |
| 105 | 16 | -92 | 18 | -5.51 | right Occipital gyrus |
| 99 | 66 | -16 | 20 | -4.95 | right Supramarginal gyrus |
| 98 | -30 | -28 | 52 | -4.83 | left Precentral gyrus |
| 74 | 34 | 28 | 2 | 5.63 | right anterior Insula |
| 73 | 40 | -60 | 50 | -4.61 | right angular gyrus |
| 54 | 22 | -32 | 72 | -5.81 | right Postcentral gyrus |
| 52 | 20 | -60 | 12 | -4.85 | right Occipital gyrus |
| 35 | 8 | -44 | 62 | -4.42 | right superior Postcentral sulcus |
| 34 | 18 | -74 | 24 | -5.00 | right posterior Angular gyrus |
| 31 | -18 | -66 | 8 | -5.02 | left Occipital gyrus |
| 31 | -36 | -16 | 16 | -4.80 | left posterior Insula |
| 29 | 54 | 4 | 36 | 5.11 | right Precentral gyrus |
| 27 | -54 | -22 | 54 | -4.95 | left Postcentral gyrus |
| 25 | -26 | -70 | -28 | -4.45 | left Cerebellum |
| 24 | -60 | -32 | 16 | -4.51 | left Supramarginal gyrus |
| 24 | 38 | -16 | 40 | -5.01 | right Postcentral gyrus |
| 23 | 56 | -58 | -6 | -4.01 | right Inferior Temporal gyrus |
| 22 | 28 | -10 | -20 | -4.16 | right Amygdala |
| 22 | -50 | 22 | -8 | -4.66 | left Inferior Temporal gyrus |
| 21 | -36 | -18 | 44 | -5.06 | left Postcentral gyrus |
| 20 | -54 | 24 | 18 | -4.34 | left Inferior Frontal gyrus |
| 20 | -50 | -20 | 18 | -4.87 | left Parietal Operculum |
| 20 | -2 | 22 | 54 | -4.24 | left Paracentral lobule |
| 19 | 32 | -76 | -38 | -4.32 | right Cerebellum |
| 18 | 58 | -58 | 28 | -4.07 | right Supramarginal gyrus |
| 17 | 44 | -70 | -20 | -3.88 | right Inferior Temporal gyrus |
| 15 | 54 | 20 | -4 | -4.59 | right Inferior Temporal gyrus |
| 15 | -8 | -76 | 16 | -4.19 | left Precuneus |
| 15 | 32 | -36 | 50 | 4.21 | right Postcentral gyrus |
| 14 | -52 | 22 | 24 | -3.98 | left Middle Frontal gyrus |
| 13 | -36 | -60 | -42 | -3.70 | left Cerebellum |

**Table 6.**
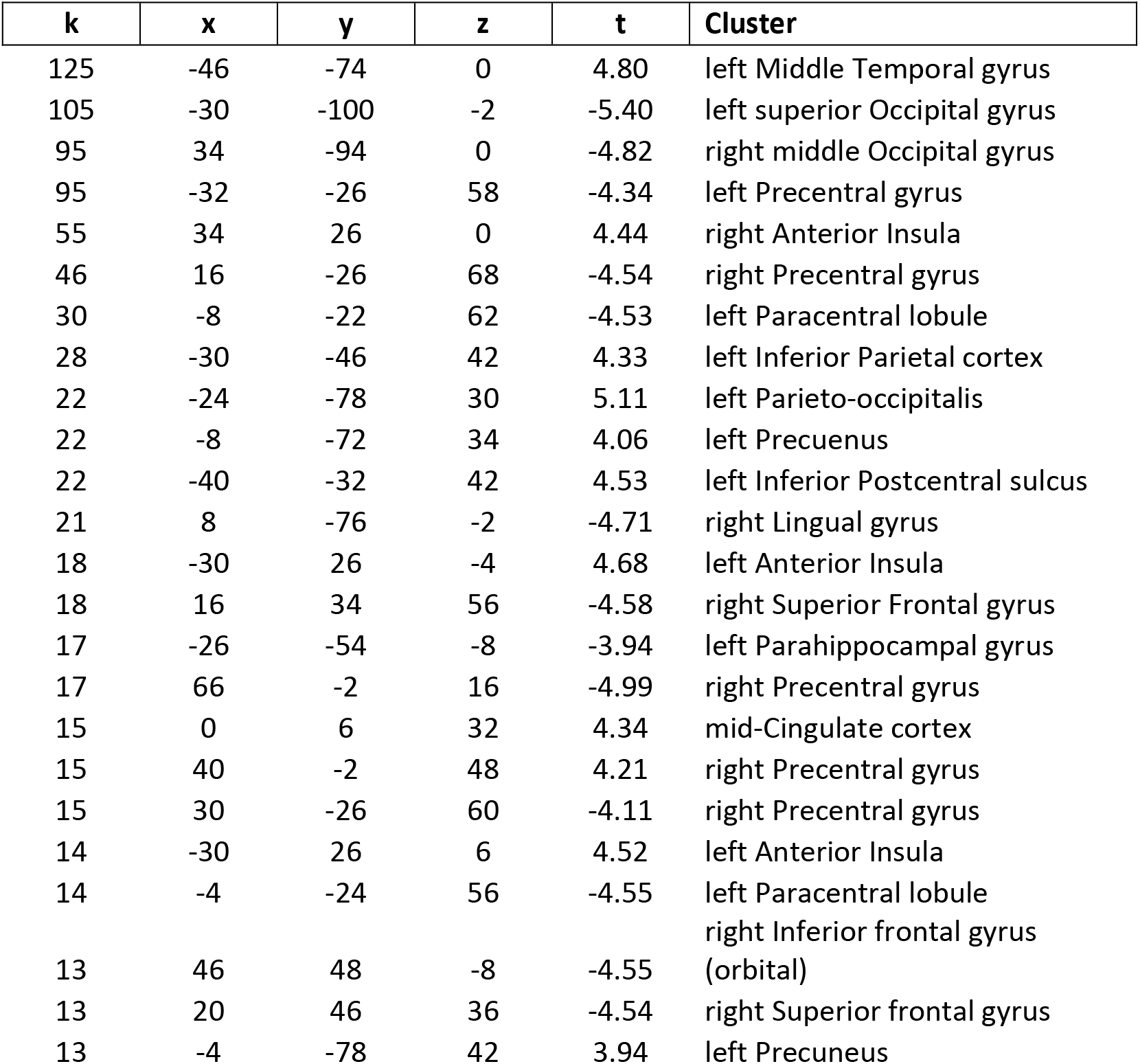
Clusters that exhibited the direction x speed interaction in voxelwise analysis at whole-brain cluster-level corrected alpha of 0.05 (peak MNI coordinates, t(84) values, and cluster size [k] refers to number of 2.0 × 2.0 × 2.0 mm^3^ voxels). Peak coordinates are presented for completeness and potential meta-analysis; with cluster-based thresholding, it is not possible to conclude that all the reported peaks were activated [see Woo et al., 2014]

**Table 7.** Clusters that exhibited the proximity x speed interaction in voxelwise analysis at whole-brain cluster-level corrected alpha of 0.05 (peak MNI coordinates, t(84) values, and cluster size [k] refers to number of 2.0 × 2.0 × 2.0 mm^3^ voxels). Peak coordinates are presented for completeness and potential meta-analysis; with cluster-based thresholding, it is not possible to conclude that all the reported peaks were activated [see Woo et al., 2014]

| **k** | **x** | **y** | **z** | **t** | **Cluster** |
| --- | --- | --- | --- | --- | --- |
| 116 | 4 | -2 | 56 | 5.58 | Supplementary Motor area |
| 50 | -6 | 16 | 34 | 4.68 | left mid-Cingulate cortex |
| 46 | 48 | -66 | 4 | -5.23 | right Middle Temporal gyrus |
| 39 | 26 | -90 | 6 | -5.35 | right Occipital gyrus |
| 37 | -30 | 28 | 2 | 5.25 | left Anterior Insula |
| 33 | 30 | -74 | 38 | -4.46 | right Parieto-occipitalis |
| 23 | 40 | -80 | 20 | -4.38 | right Occipital gyrus |
| 22 | 8 | 22 | 38 | 4.65 | right mid-Cingulate cortex |
| 21 | -62 | 4 | 8 | 4.19 | left Precentral gyrus |
| 19 | 18 | -66 | 52 | -4.47 | right Superior Parietal lobule |
| 15 | 46 | -58 | -6 | -4.49 | right Inferior Temporal gyrus |
| 13 | -24 | -96 | 2 | -4.09 | left middle Occipital gyrus |
| 13 | 12 | -72 | 6 | 4.15 | right Occipital gyrus |
| 13 | 14 | -54 | 64 | -3.85 | right Superior Parietal lobule |

**Table 8.** Clusters that exhibited the proximity x direction x speed interaction in voxelwise analysis at whole-brain cluster-level corrected alpha of 0.05 (peak MNI coordinates, t(84) values, and cluster size [k] refers to number of 2.0 × 2.0 × 2.0 mm^3^ voxels). Peak coordinates are presented for completeness and potential meta-analysis; with cluster-based thresholding, it is not possible to conclude that all the reported peaks were activated [see Woo et al., 2014]

| **k** | **x** | **y** | **z** | **t** | **Cluster** |
| --- | --- | --- | --- | --- | --- |
| 17 | -54 | -24 | 38 | 3.98 | left Central sulcus |
| 17 | 24 | 30 | 52 | -5.02 | right Superior Frontal gyrus |

### BST ROI analysis

Given that the BST is a rather small region that is involved in threat-related processing, we ran a focused ROI analysis using anatomically defined left/right BST masks, and unsmoothed data to minimize the influence of signals from surrounding structures. We observed a robust effect of threat proximity in the right BST (and weak evidence in the left BST), with stronger responses when circles were closer than farther (Figure 9A; Table 9). For the right BST, some evidence for proximity by speed interaction was seen.

**Figure 9.**
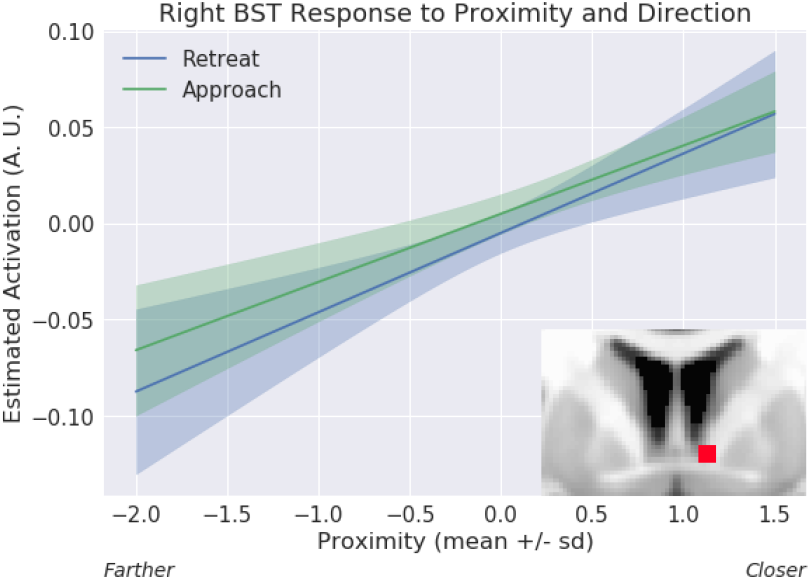
Proximity effect in the bed nucleus of the stria terminalis (BST) ROI analysis. Estimated responses for a range of proximity values. Activity increased as a function of proximity for both approach and retreat. The confidence bands were obtained by considering variability during approach and retreat, separately; see Methods. A.U.: arbitrary units.

**Table 9.** BST ROI analysis results.

| Regressor | Left BST |  | Right BST |  |
| --- | --- | --- | --- | --- |
|  | t(84) | p-value | t(84) | p-value |
| Proximity | 1.91 | 0.0591 | 4.17 | 0.0001 |
| Direction | 1.29 | 0.1997 | 0.64 | 0.5263 |
| Speed | 0.83 | 0.4099 | 1.78 | 0.0780 |
| DirectionXSpeed | -0.91 | 0.3634 | -0.26 | 0.7940 |
| ProximityXDirection | -1.63 | 0.1066 | -0.35 | 0.7268 |
| ProximityXSpeed | -0.08 | 0.9353 | 2.60 | 0.0109 |
| ProximityXDirectionXSpeed | 0.00 | 0.9986 | -1.69 | 0.0956 |

**Bonferroni correction for multiple comparisons: $0.05/7 = 0.0071$**

### Relationship between SCR responses and brain activity

We evaluated the linear relationship between SCR and fMRI by running a robust correlation analysis (across subjects). Because multiple aspects of both the SCR and fMRI data could be probed (simple effects and interactions), we chose to focus the interrogation on the proximity by direction interaction. Thus, for both SCR and fMRI, the strength of the two-way interaction was considered for the analysis (as given by the regression coefficient in equation 1). To minimize the problem of multiple statistical comparisons, for this analysis, we focused on clusters exhibiting a two-way interaction in the right anterior insula and the right amygdala, regions that feature in most models of threat processing. We did not detect a relationship between SCR and fMRI responses in either the right anterior insula (r(77) = 0.07, P = 0.550) or the right amygdala (r(75) = −0.04, P = 0.697).

### Relationship between anticipatory activity and physical shock responses

Our interpretation of the proximity by direction interaction was that it reflected, at least in part, threat-related processing, especially in brain regions important for this type of processing, such as the anterior insula. In an exploratory analysis, we tested if the strength of this interaction effect was associated (across participants) with the strength of responses evoked by physical shock. For the right anterior insula cluster that exhibited a proximity by direction interaction, we detected a positive linear relationship between the two measures (r(80) = 0.33, P = 0.002; Figure 10). Given the importance of the amygdala in threat processing, we also tested the relationship in the right amygdala (also considering the cluster that exhibited a proximity by direction interaction), but no effect was detected (r(80) = −0.02, P = 0.888).

**Figure 10.**
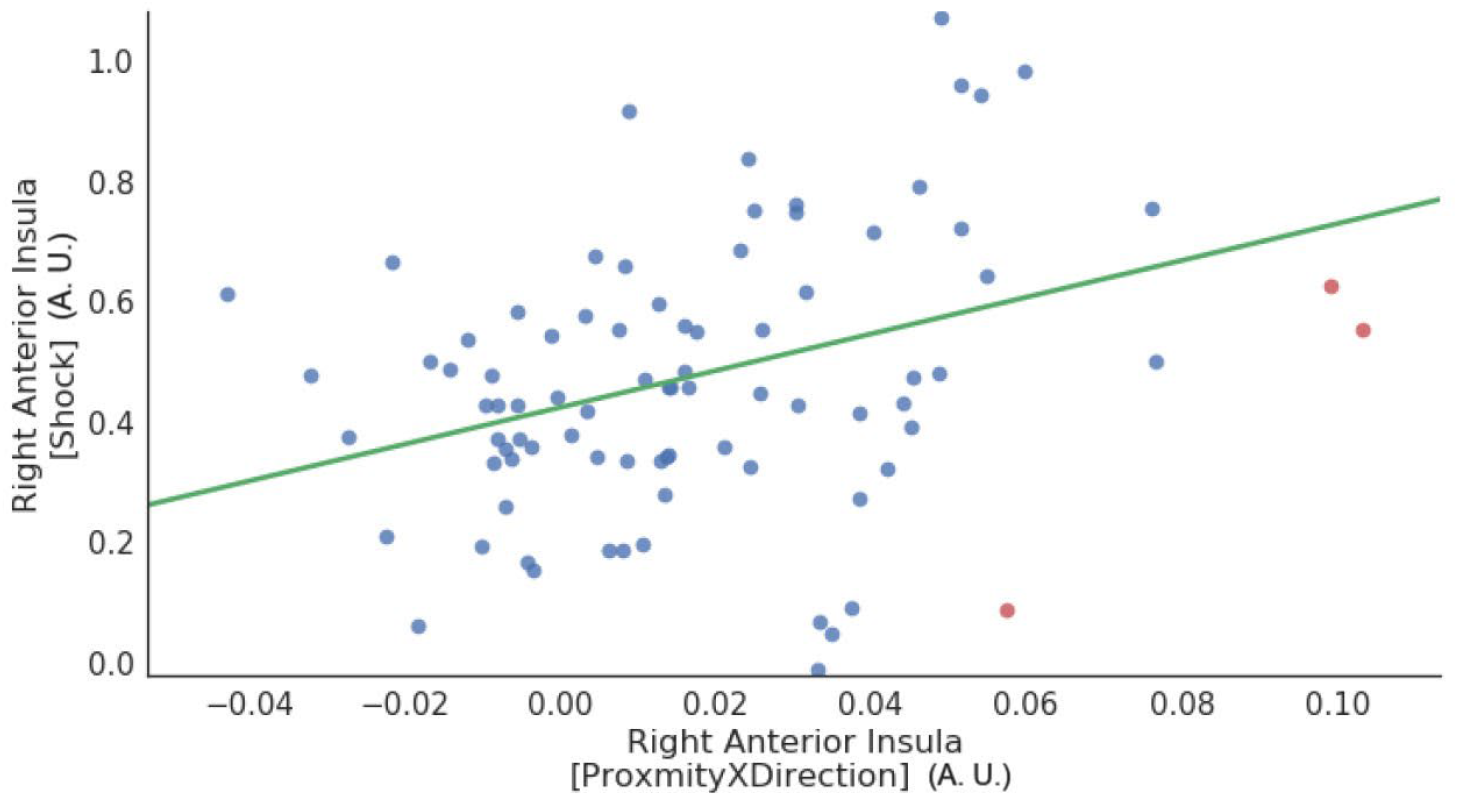
Relationship between anticipatory activity and physical shock responses in the right anterior insula. For the anticipatory activity, the proximity by direction interaction was considered for the analysis. Data points correspond to participants (red points indicate outliers deemed based on the robust correlation algorithm). A.U.: arbitrary units.

### Individual differences in state and trait anxiety

Linear relationships between state/trait anxiety and SCR or, separately, fMRI interactions of proximity and direction in the right anterior insula were not detected (all rs < 0.1 in absolute value). We detected a modest positive relationship between state anxiety and fMRI interactions of proximity and direction in the right amygdala (in the cluster that exhibited a proximity by direction interaction; state: r(77) = 0.2107, p-value = 0.0544; trait: r(79) = 0.0769, p-value = 0.4870). Given the multiple tests involved here, we do not believe these findings are noteworthy.

### Exploratory analyses: PAG responses

To visualize the responses of the PAG, we plotted estimated responses (Figure 11A), as done above for the right anterior insula, right amygdala, and right BST. To do so, we employed the cluster (38 voxels) that exhibited the direction effect (approach vs. retreat) previously reported (Figure 6). Upon plotting, we discerned an effect of proximity for the approach condition, but not for retreat, consistent with a proximity by direction interaction (which was not detected in the voxelwise analysis). Given the importance of the PAG in the orchestration of defensive responses in the face of threat (Bandler and Shipley, 1994; Pessoa, 2016), we performed an additional exploratory analysis in this region. First, we generated a representative time series for the PAG by averaging the time series of the voxels within the cluster (based on the voxelwise effect of direction), and then evaluated the full model (Equation 1). As shown in Table 10, a robust proximity by direction interaction was detected (note that the interaction effects were nearly independent from the selection criterion, which was based on direction; the correlation between the interaction and direction was −0.14). Given this result, we inspected again the results at the voxelwise level, and observed some voxels that exhibited such an interaction, but too few to survive cluster thresholding.

**Figure 11.**
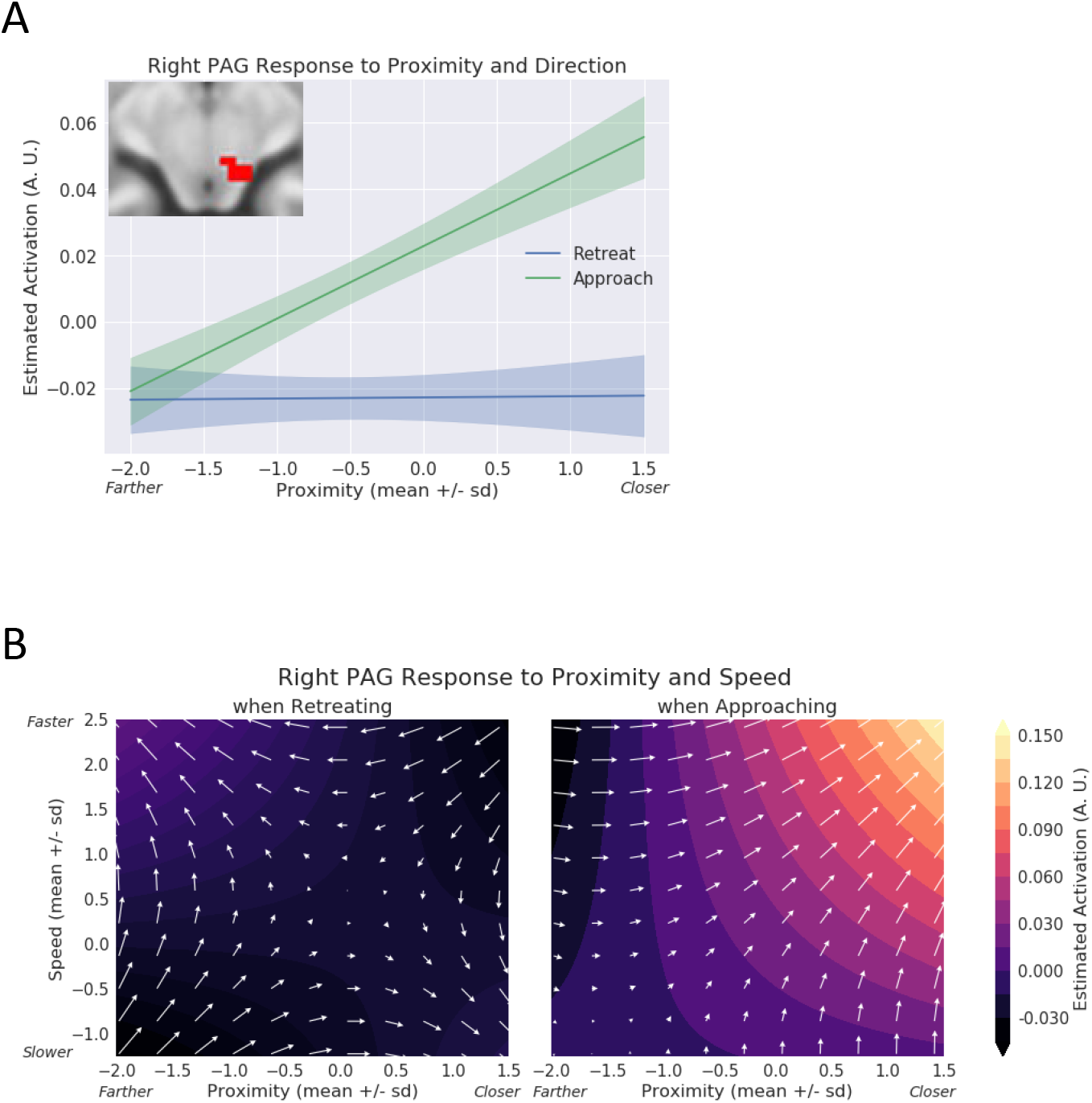
Exploratory analysis of the periaqueductal gray (PAG). (A) Estimated responses for a range of proximity values. During approach, activity increased as a function of proximity; activity changed little during retreat periods. The confidence bands were obtained by considering within-subject differences (approach minus retreat); see Methods. A.U.: arbitrary units. (B) Contour plots show estimated responses for different combinations of proximity and speed during approach and retreat periods. Arrows point in the direction of signal increase. During approach, both proximity and speed simultaneously influenced responses, which increased when the circles were closer and speed was higher. A.U.: arbitrary units.

**Table 10.** Exploratory right PAG ROI analysis results.

| Regressor | t(84) | p-value |
| --- | --- | --- |
| Proximity | 2.33 | 0.0220 |
| Direction | 6.75 | 0.0000 |
| Speed | 2.69 | 0.0087 |
| DirectionXSpeed | 1.92 | 0.0588 |
| ProximityXDirection | 3.89 | 0.0002 |
| ProximityXSpeed | 1.17 | 0.2455 |
| ProximityXDirectionXSpeed | 2.86 | 0.0054 |

**Bonferroni correction for multiple comparisons: $0.05/7 = 0.0071$**

Notably, we also observed a robust three-way interaction. As the three factors simultaneously affected PAG responses, the finding can be visualized via a contour plot (Figure 11B). During approach periods, when proximity increased (circles moved closer to each other), stronger responses were observed as speed increased from slower to faster (compare the upper-right vs. lower-left quadrants).

### Exploratory analyses: Potential nonlinear effects of proximity

The regression model we employed (equation 1) makes the assumption that the effect of proximity is linear. In additional exploratory analyses, we investigated potential nonlinear effects of proximity on brain activity. To do so, we inspected the pattern of the residuals as a function of proximity in the right anterior insula, right amygdala, right BST, and right PAG. For example, Figure 12 shows the residuals when employing equation 1 for the right anterior insula. Based on the pattern of residuals, the linear modeling approach adopted here appears to be reasonable in the context of our experiment.

**Figure 12.**
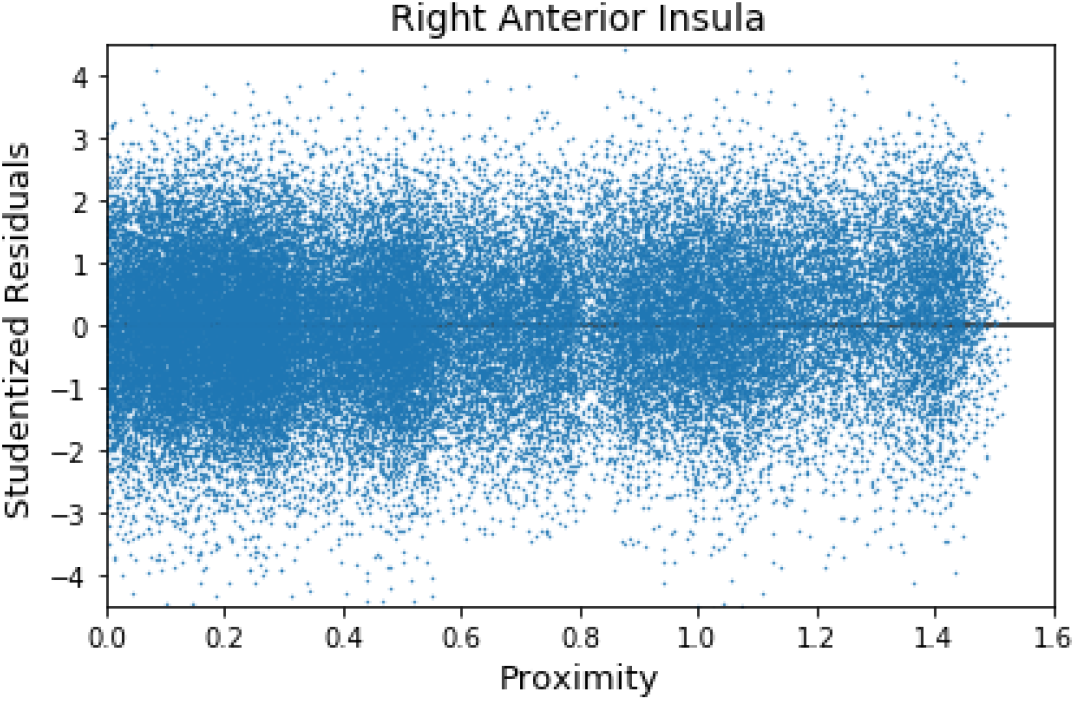
Exploratory analysis of potential nonlinear effects of proximity. The residuals from the model fit are plotted as a function of proximity. No appreciable lack of fit is evident. To plot residuals for all participants, they were first studentized (jitter as a function of proximity was also used to reduce overlap).

## Discussion

In the present study, we investigated the role of threat-related factors and their temporally evolving interactions. Our findings support the view that threat processing is context sensitive and dynamic (Fanselow and Lester, 1988; Blanchard and Blanchard, 1990; Kavaliers and Choleris, 2001; Mobbs et al., 2015). In some brain regions, signal fluctuations were sensitive to continuous manipulations of proximity and speed indicating that threat processing is dynamic. Importantly, whereas some brain regions tracked individual threat-related factors (proximity, direction, or speed), others were also sensitive to combinations of these variables revealing the context-sensitive nature of threat processing. In this section, we will focus the discussion on a few of the brain regions that have been most heavily implicated in threat-related processing in the literature, specifically the anterior insula, amygdala, BST, and PAG.

To investigate how threat-related factors influence physiological arousal during dynamic threat anticipation, we recorded SCR during scanning. We observed robust effects of proximity and direction, with larger responses during near vs. far and approach vs. retreat, respectively. Of note, we observed a robust proximity by direction interaction, where responses to threat direction (approach vs. retreat) were enhanced when the circles were *near* compared to far suggesting that the influence of dynamic threat anticipation on physiological arousal was context dependent.

Responses in the anterior insula were driven by proximity, direction, and speed. Importantly, in the right hemisphere, anterior insula responses also exhibited an interaction between proximity and direction, such that the approach vs. retreat contrast was enhanced when the circles were *near* compared to far. The anterior insula supports subjective awareness of bodily states (Craig, 2002, 2009), and is consistently engaged during threat-related processing (Nitschke et al., 2006; Simmons et al., 2006). In particular, the anterior insula is implicated in tracking threat proximity and direction during aversive anticipation (Mobbs et al., 2010; Somerville et al., 2010). Our results replicated these findings while extending them by showing that the effects of threat proximity and direction are not independent but *jointly* contribute to responses in the anterior insula.

In the present study, we observed a proximity by direction interaction in the right amygdala, but in the opposite direction to that seen in the anterior insula: when far, proximity had a weak or no effect on responses, but when near responses were greater for retreat relative to approach. In fact, the differential response to retreat vs. approach became more pronounced as the circles approached each other, with approach responses decreasing with increased proximity. In paradigms investigating the independent effects of proximity and direction on threat anticipation, Somerville and colleagues (2010) suggested a limited role of the amygdala in tracking threat proximity, whereas Mobbs and colleagues (2010) observed amygdala responses that responded to the proximity and direction of threat. In a study involving virtual predators, Mobbs and colleagues (2007) reported increased activation in the dorsal amygdala when threat was near, whereas responses were *stronger* in the inferior-lateral amygdala with distant threats. Thus, our results more closely resemble the latter amygdala sub-region. It should be noted that in previous studies, similar to the pattern of responses observed in the current study, we and others have observed amygdala *deactivations* during short and long periods of sustained threat (relative to safe conditions; Choi et al., 2012; McMenamin et al., 2014; Grupe et al., 2016); see also (Pruessner et al., 2008; Wager et al., 2009) in case of social stress/threat.

The role of the BST in threat processing has gained increased attention in the past two decades (Davis and Whalen, 2001; Shackman and Fox, 2016), especially during conditions involving temporally extended and less predictable threats. Given the small size of the structure and its anatomical location, studying the BST with fMRI is particularly challenging. Recently, anatomical masks for both regular and higher field scanning have been published (Avery et al., 2014; Torrisi et al., 2015; Theiss et al., 2017), which should enhance the reproducibility of published findings. We analyzed BST data using an anatomical mask and unsmoothed data, which is important because nearly all studies have employed some voxelwise spatial smoothing which blends BST signals with those of adjacent territories (beyond the inherent point spread function of imaging itself); but note that smoothing within the BST was accomplished by averaging unsmoothed time series of voxels within the anatomically defined ROI. In the right BST, we observed an effect of proximity, and a proximity by speed interaction (but note that these effects were less robust as they would not survive correction for the 7 tests employed; or 14 if one were to consider both hemispheres). The observed effect of proximity is consistent with previous findings that the BST responds to threat proximity (independent of direction; Somerville et al., 2010; Mobbs et al., 2010; although the activated region was sufficiently large as to make anatomical localization challenging in the study by Mobbs and colleagues).

The PAG of the midbrain has been implicated in aversive and defensive reactions (Bandler, 1988; Bandler and Shipley, 1994), in line with more recent studies (Tovote et al., 2016). In humans, the PAG has been suggested to be involved in negative emotional processing more generally (Lindquist et al., 2012; Satpute et al., 2013). The virtual tarantula manipulation by Mobbs and colleagues (2010), where participants were shown a prerecorded video of a spider moving towards or away from their feet was particularly effective in engaging the PAG when threat was proximal (although the activation was very extensive, and thus difficult to localize). Here, in the voxelwise analysis, we only detected an effect of direction in the right midbrain/PAG where stronger responses were observed when circles were approaching compared to retreating. However, exploratory analyses revealed a robust proximity by direction interaction, as well as a proximity by direction by speed interaction. These results are potentially important because they suggest that threat-related responses in the PAG are sensitive to multiple factors that jointly determine the PAG’s activity. Interestingly, unlike in the amygdala and anterior insula where we only observed an interaction between proximity and direction, speed also played a role in the PAG. However, given the exploratory nature of our analysis, future converging findings are needed to more precisely delineate the role of multiple threat-related factors on PAG activity during aversive anticipation.

A limitation of the present study was that it did not include two types of control condition. First, only aversive events were encountered and not motivationally positive ones. Thus, the extent to which signals investigated here were linked to threat and not “motivational significance” more generally needs to be further investigated. Second, because a “no-shock condition” was not included, it is possible that signal fluctuations were due to processes linked to tracking circle movement, including predicting future circle positions based on current position and prior movement statistics. In this context, the anterior insula is an interesting case because it is a highly functionally diverse region and is sensitive to a very broad range of influences (Anderson et al., 2013). But because anterior insula signals were sensitive to *interactions* between proximity and direction, it is unlikely that prediction/updating processes explained responses, as participants presumably engaged in such processing in a similar fashion when the circles were closer or father. In addition, we observed a positive correlation between proximity by direction interaction responses and responses evoked to physical shock, consistent with the fact that responses were at least in part related to anticipation of the aversive event. Finally, and more generally, the three-way interaction in the PAG (but see Results for the exploratory aspect of this result) exhibited a degree of specificity (compare left and right panels in Figure 11B) that are difficult to explain by visuo-cognitive processes of circle movement tracking.

Another limitation of the preset study was that participants did not have control over the threat. Unlike active avoidance paradigms where participants could perform instrumental actions to terminate or completely avoid the threat (for instance, see Mobbs et al., 2007), the passive nature of our task likely constrained the types of “defensive processing” observed. In particular, investigation of a richer set of behaviors and brain responses, such as described in the threat imminence continuum framework (Fanselow and Lester, 1988), will require novel approaches and experimental designs attuned to findings in ethology and behavioral ecology (see Mobbs et al., 2018).

To conclude, we investigated how multiple threat-related factors (proximity, direction, and speed) interact when varied continuously. In particular, we asked whether signal fluctuations in brain regions track threat-related factors dynamically? If so, to what factor(s) and factor combinations are they sensitive? We observed a proximity by direction interaction in the anterior insula where approach vs. retreat responses were enhanced when threat was proximal. In the right amygdala, we also observed a proximity by direction interaction, but in the opposite direction as that found for the anterior insula; retreat responses were stronger than approach responses when threat was proximal. In the right BST, we observed an effect of proximity and in the right PAG/midbrain we observed an effect of direction as well as a proximity by direction by speed interaction (the latter was detected in exploratory analyses but not in a voxelwise fashion). Overall, this study refines our understanding of the mechanisms involved during aversive anticipation in the typical human brain. Importantly, it emphasizes that threat processing should be understood in a manner that is both context sensitive and dynamic. As aberrations in aversive anticipation are believed to play a major role in disorders such as anxiety and depression (Grupe and Nitschke, 2013; Dillon et al., 2014), our findings of interactions between multiple threat-related factors in regions such as the amygdala, anterior insula, and PAG may inform the understanding of brain mechanisms that are dysregulated in these disorders.

## Acknowledgements

We would like to thank Brenton McMenamin for paradigm development, Dan Levitas for data collection, Jason Smith for help with processing scripts, Mahshid Najafi for help with initial pre-processing of the fMRI data, and Nicole Friedman and Jessica Berman for help with subject recruitment. The authors also acknowledge the Behavioral and Social Sciences College, University of Maryland, high performance computing resources (http://bsos.umd.edu/oacs/bsos-high-performance) made available for conducting the research reported in this paper.

